# Cryptic diversity in *Astyanax* (Characiformes: Acestrorhamphidae) from the Magdalena basin, Colombia: Insights from molecular and morphometric evidence

**DOI:** 10.64898/2026.03.28.714954

**Authors:** Edna Judith Márquez, Kevin León García-Castro, Daniela Ramírez Álvarez, Carlos DoNascimiento

**Author notes:** Corresponding author (EJM).

## Abstract

*Astyanax* Baird & Girard, 1854 is a widely distributed and species-rich genus of Acestrorhamphidae, whose abundant populations in Neotropical basins play a crucial ecological role at the trophic level. Taxonomic uncertainties persist within the genus, as seen in *Astyanax* sp. (formerly designated as *A. fasciatus*) from the Magdalena basin in Colombia. Concerns about its genetic status are heightened due to ecological threats posed by hydroelectric dams, from habitat loss to river connectivity. We isolated and characterized 17 microsatellite loci to assess the population genetics of this species in a broad sample from the middle and lower sections of the Cauca River, now interrupted by the Ituango dam. Furthermore, a multidisciplinary approach integrating phylogenetic analyses of mitochondrial (*COI*) and nuclear (*rag2*) markers with geometric morphometric analyses was employed to evaluate potential cryptic diversity within *Astyanax* sp. Microsatellites revealed two genetic groups in the studied area, strongly supported as distinct lineages by phylogenetic analyses. Unexpectedly, one of these lineages of *Astyanax* sp. was recovered in an unresolved clade with samples of *A. microlepis* and allopatric samples of *A. viejita* from the Maracaibo Lake basin. Each genetic group showed high genetic diversity, but also evidence of recent bottleneck events and significant-high values of inbreeding. Morphometric analyses provided evidence of significant phenotypic differentiation among *A. microlepis*, *Astyanax* sp. 1 (Asp1), and *Astyanax* sp. 2 (Asp2). Morphological patterns ranged from the robust profile of *A. microlepis* to the streamlined shape of *Astyanax* sp. 2 (Asp2), with *Astyanax* sp. 1 (Asp1) displaying intermediate traits and localized differences in head length and fin placement. Statistical support from permutation tests and a high overall classification accuracy (95.65%) underscore the existence of distinct morphospecies, suggesting that phenotypic differentiation is well-established, despite the complex evolutionary history of the group. This study suggests the presence of cryptic diversity within *Astyanax* sp. and provides valuable genetic information for the conservation and management of their populations in the Magdalena basin.

## Introduction

*Astyanax* Baird & Girard, 1854 is a genus of tetra fishes with broad distribution across the American continent, extending from Texas to Patagonia, and is especially abundant in the Amazon and Orinoco basins. The genus currently includes 122 valid species, comprising approximately 2.8% of the taxonomic diversity of the entire order Characiformes [1], and recently was reallocated into the family Acestrorhamphidae from its traditional placement in Characidae [2]. Due to non-exclusive diagnostic characters, phenotypic plasticity, and limited resolution by *COI*-based molecular analyses, *Astyanax* remains a non-monophyletic group [2–5], complicating species-level taxonomic identification within the genus.

In Colombia, 22 *Astyanax* species are reported, nine of them in the Magdalena basin [6], including *Astyanax* sp., formerly designated as *Astyanax fasciatus* (Cuvier, 1819) [7]. *Astyanax fasciatus* was reallocated in *Psalidodon* Eigenmann, 1911 [5] and its geographic distribution restricted to the São Francisco basin in southeastern Brazil [5], leaving at least five allopatric species from remote cis-Andean or even trans-Andean basins that are still listed as synonyms of *P. fasciatus* [8], in a taxonomic limbo.

Species of *Astyanax* from the Magdalena basin have been reported as omnivorous, with seasonal life strategies, early maturation, body size ranging from 60 to 183 mm of standard length (SL), and short-range migrations [9]. They are also considered the most abundant species both in numerical abundance and biomass in the Magdalena basin, even in reservoir systems, and play a fundamental role in the food chain [9]. Local fishermen recognize that *Astyanax* species are among the first populations to leave floodplains and signal upstream migrations for other species during periods of water descent [9,10]. Some of these species, such as *A. magdalenae* Eigenmann & Henn, 1916 and *Astyanax* sp., have also been reported for ornamental and/or consumption purposes in the last two decades [9].

Despite *Astyanax’s* high species diversity, taxonomic uncertainty, and multiple threats, few studies have examined its microevolutionary genetics scale. These studies have focused mainly on populations from Mexico and Brazil [11–14]). In Colombia, only one study has assessed population genetics of *A. caucanus* (Steindachner, 1879), using species-specific microsatellite loci [15]. That study reported high gene flow along a 340 km stretch of the Cauca River, even among sectors recently isolated by the Ituango dam, as well as high genetic diversity (as in other congeners) and large effective population sizes [15]. However, it also found evidence of recent bottleneck events and significant inbreeding [15].

*Astyanax* is characterized by high phenotypic plasticity and a remarkable capacity to adapt to diverse habitats [16,17]. This plasticity has motivated numerous studies using geometric morphometrics, a tool for quantifying body shape variation among genera, species, populations, and sexes [18]. Such data are crucial for refining species boundaries and assessing the influence of both genetic and environmental factors [19–23], especially in groups like Characidae, where morphological conservatism often masks underlying genetic differences.

Several studies have combined genetics and morphometrics to explore their interplay in *Astyanax* and related genera [23–27]. For instance, ecologically distinct morphs of *Astyanax* have been identified in the San Juan River basin (Central America), with marked morphological differentiation (one morph with elongated body and predatory features and the other morph with deep body and generalist traits), despite limited genetic divergence between them [22]. Morphological distinction has also been corroborated by genetic evidence in the *Psalidodon scabripinnis* (Jenyns, 1842) complex, where *P. paranae* (Eigenmann, 1914) exhibits a more fusiform body than *P. rivularis* (Lütken, 1875) [26]. However, a clear correspondence between morphological and genetic patterns is often absent. Studies on sympatric populations of *A. caballeroi* (Contreras-Balderas & Rivera-Teillery, 1985) and *A. aeneus* (Günther, 1860) in Mexico have revealed significant differences in body depth, head profile, and fin morphology that are not reflected by the molecular data [19].

Geometric morphometrics also enables analyses of environmental effects on body shape, such as habitat type and water flow [28–31]. Ecomorphological divergence is well-documented in *Astyanax* and related genera. A common pattern is the correlation between body shape and water velocity, species inhabiting lotic environments, such as *Psalidodon rivularis* and *P. paranae*, typically have fusiform bodies adapted to swimming against currents, whereas species from lentic habitats, like *A. abramis* (Jenyns, 1842), *A. lacustris* (Lütken, 1875), and *A. asuncionensis* Géry, 1972, exhibit deeper, laterally compressed bodies, suited for maneuverability [32]. This pattern also occurs within single species. In river populations, *A. lacustris* have a fusiform shape with a deep caudal peduncle, while their lagoon-dwelling counterparts are deeper-bodied with a shallower caudal peduncle [33]. Likewise, *A. bimaculatus* (Linnaeus, 1758) from reservoirs have deeper bodies and longer fin bases than those from rivers [32]. Some species that occupy both environments, such as *P. fasciatus*, display an intermediate body shape, reflecting their ecological flexibility [23,32].

In the present work, we designed and evaluated a set of microsatellite loci to assess genetic diversity and population structure of *Astyanax* sp. in the middle and lower sections of the Cauca River, spanning sectors upstream and downstream of the Ituango dam. Additionally, *COI* and *rag2* sequences were compared within a broad phylogenetic framework (with emphasis on trans-Andean representatives of *Astyanax* and species of the resurrected *Psalidodon* [5], including the true nominal *P. fasciatus*), to determine whether groups potentially identified through microsatellites correspond to divergent lineages within the genus. Finally, geometric morphometric analyses were conducted to investigate potential drivers of diversification in *Astyanax*.

## Materials and methods

### Sampling

The Cauca River, stretching 1,350 km from Laguna del Buey in the south to its confluence with the Magdalena River, flows along the inter-Andean valley and canyon delimited by the Western and Central Cordilleras of the northern Andes mountains, with its upper section at around 900 m asl, the final 500 km features a steep, narrow canyon, dominated by rapids and waterfalls, which posed challenges for migrating fishes, even before the Ituango dam construction. This dam is the largest in Colombia, standing at 225 m high, and it is expected to contribute approximately 2,400 MW of power annually [34,35].

Between 2019 and 2021, a total of 324 muscle and fin tissue samples of *Astyanax* sp. were collected from eight sections in the middle and lower parts of the Cauca River (**Fig 1**). High-resolution photographs were captured for each specimen and standardized for landmark digitization, providing the framework for subsequent geometric morphometric analyses. These sections were chosen due to their contrasting hydrological and geographical characteristics [36]. Two sections, namely S1 and PHI, were located upstream of the Ituango dam, while the remaining sections (S2, S3, S4, S5, S6, and S7/S8) were situated downstream of the dam. The PHI section represents the current reservoir of the dam. Collections were conducted by Grupo de Ictiología from the Universidad de Antioquia (GIUA), Fundación Humedales, and Grupo de Biotecnología Animal from Universidad Nacional de Colombia, campus Medellín.

**Fig 1.**
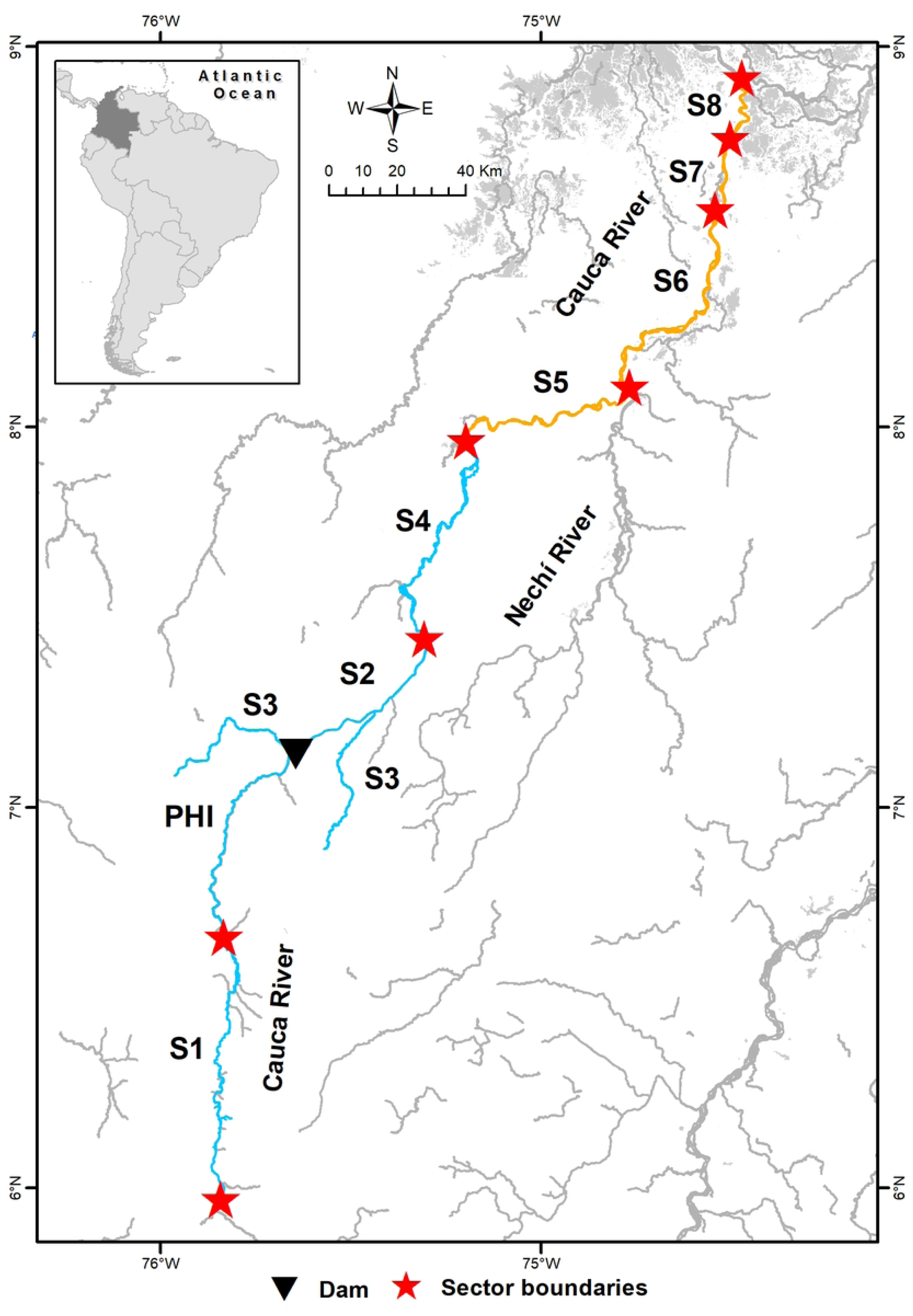
Geographic location of sampling sectors in the Cauca River basin, Colombia. Blue lines represent upstream and dam-proximate sectors (S1, PHI, S2, S3, and S4) predominated by genetic stock Asp1, while orange lines denote downstream sectors (S5, S6, S7, and S8) predominated by genetic stock Asp2 (see Results). The map was generated by the authors based on 1:100,000 scale cartographic data from the Instituto Geográfico Agustín Codazzi (IGAC, 2019). Data available at: https://geoportal.igac.gov.co/.

### Isolation and characterization of microsatellite loci

We designed and assessed 42 specific primer pairs (Afa01-Afa42; S1 Table) to amplify microsatellite loci containing tri-, tetra-, and pentanucleotide motifs to assess population genetics in samples of *Astyanax* sp. We applied Illumina’s MiSeq v2 technology and followed a previously established methodology [36] to identify and isolate microsatellite loci, using a sample (P730) of *Astyanax* sp. collected in the Cauca River. Prinseqlite v0.20.4 was used to refine the 250-base paired-end reads generated in the sequencing, followed by Abyss v1.3.5 assembly and PAL_FINDER v.0.02.03 analysis to locate microsatellite loci with perfect repeats [37]. Primer pairs for amplification were designed via Primer3 v.2.0, and electronic PCR validation ensured accurate primer alignment for the amplifiable loci.

PCR amplifications were conducted in a final reaction volume of 10 µl, following a three-primer methodology [38,39]. The reaction mixture contained 1X Platinum™ Multiplex PCR Master Mix, 0.25% Enhancer/GC, 0.20 pmol/µl of forward primer with a universal tail, 0.40 pmol/µl of reverse primer, and 0.20 pmol/µl of fluorescently labeled tail. The thermal profile involved an initial step at 95°C for 3 minutes, followed by 35 cycles of 90°C for 30 seconds and 56°C for 35 seconds, without final extension. Fragment separation was carried out on a SeqStudio Genetic Analyzer, using LIZ600 as an internal molecular marker. Allelic assignment was performed using GeneMarker v3.0.0, while Micro-Checker v.2.2.3 [40] was employed to identify potential genotyping errors in the data matrix.

We employed 36 random samples representing all sections of the studied river to characterize each of the amplified loci. These samples were used to calculate various genetic diversity metrics (detailed below), including polymorphic information content (PIC), using CERVUS v3.0.7 [41].

### Population genetics analyses

Genetic differentiation and structure among sections were assessed using three complementary approaches: (i) Bayesian analysis of co-ancestry performed in Structure v2.3.4 [42], with optimal number of populations determined via estimators in Structure Selector [43]; (ii) analysis of molecular variance (AMOVA) and the calculation of pairwise F’_ST_ and Jost’s D_EST_ indices using GenAlEx v6.502 [44]; and (iii) discriminant analysis of principal components (DAPC) implemented with the Adegenet package [45] in R v3.4.0.

To assess genetic diversity and demographic events, we calculated genetic diversity metrics (Na: average number of alleles per locus, H_O_, H_E_: observed and expected heterozygosity, respectively) and the inbreeding coefficient (F_IS_) for each section and genetic group, using Arlequin v3.5.2.2 [46]. Additionally, allelic richness (Ar) was determined using FSTAT v.2.9 [47]. Deviations from Hardy-Weinberg equilibrium (HWE) were analyzed using the web version of GENEPOP v4.7.5 [48]. The effective population size (N_e_) was estimated through NeEstimator v2.1 [49]. Furthermore, identification of genetic evidence for recent bottlenecks was assessed using the excess heterozygosity method in BOTTLENECK v1.2.02 [50] and the calculation of the M index [51]. Statistical significance in multiple comparisons was evaluated after applying the Bonferroni sequential correction.

### Phylogenetic analyses

Our study involved the amplification and concatenation of the *COI* and *rag2* genes, including samples of *Astyanax* sp. genotyped in this study and selected, according to genetic stocks detected by genetic structure analyses (Table 1). Additional sequences analyzed included *Astyanax* sp. from the Cauca and Magdalena Rivers, as well as *A. microlepis* collected at its type locality in the upper Cauca River. Furthermore, GenBank sequences of *A. caucanus*, *A. viejita sensu* Rossini *et al*., 2016 [3], from the Maracaibo Lake basin and other congeners were incorporated (Table 2). Representatives of the genera *Astyanacinus*, *Carlana*, *Ctenobrycon*, *Jupiaba*, *Oligosarcus*, *Parastremma*, and *Psalidodon* were included as outgroups (Table 2). Taxonomic nomenclature of species names followed [2].

**Table 1.**
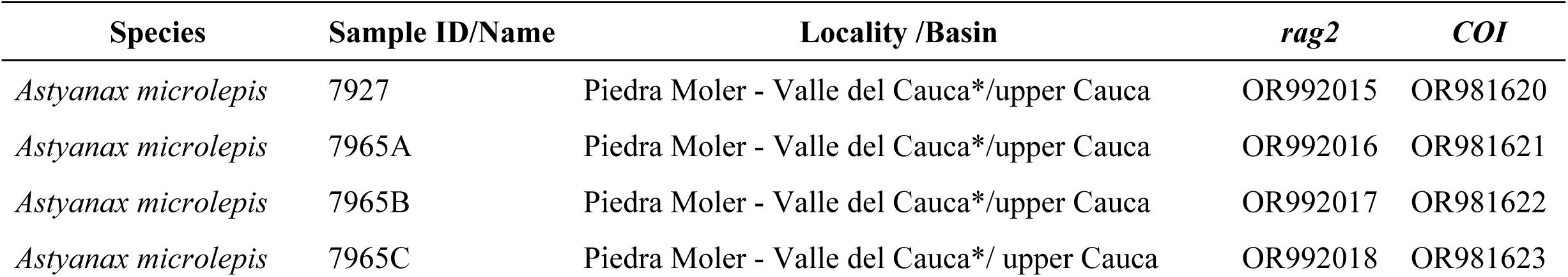
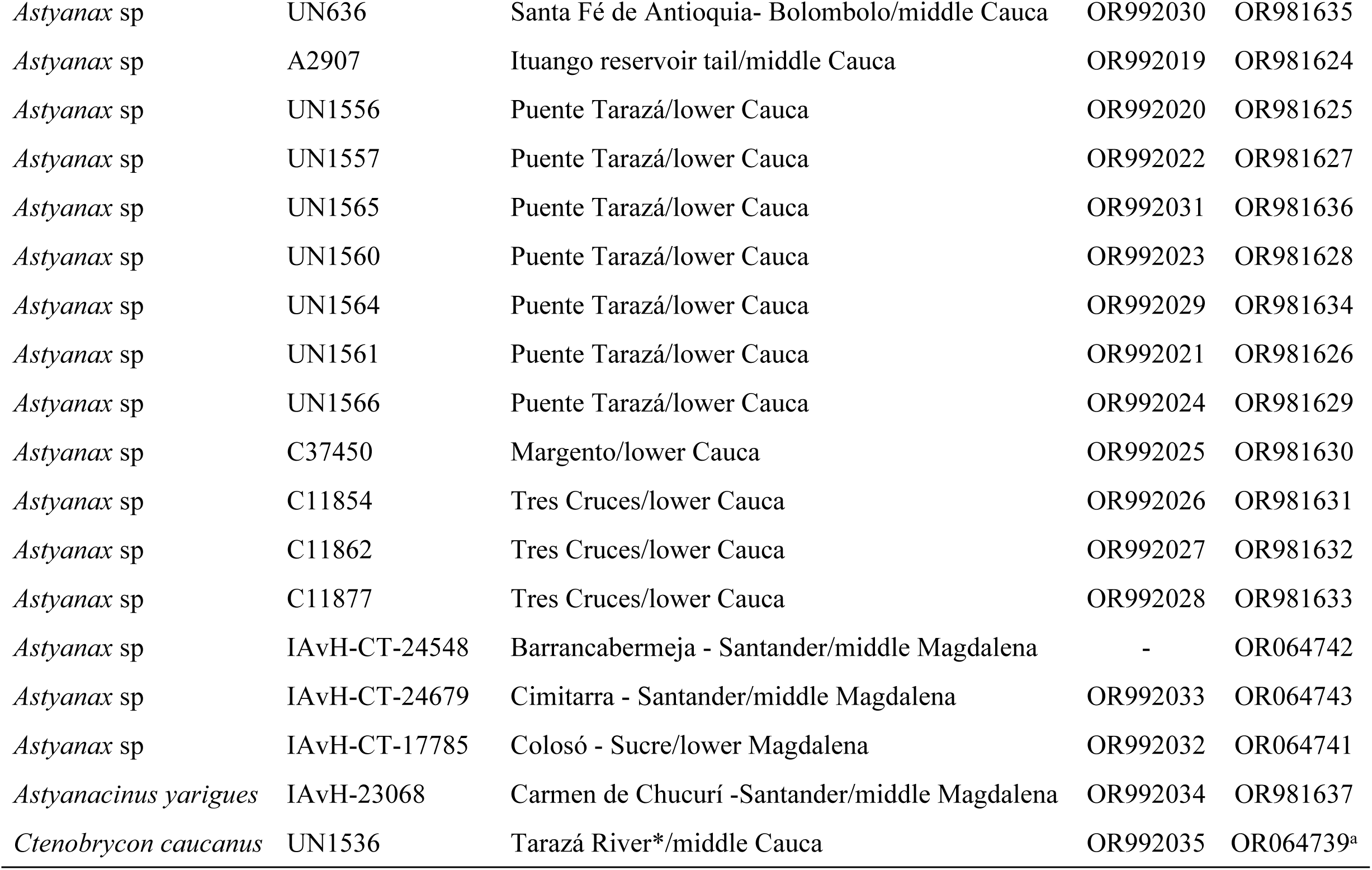
List of sequences of *Astyanax*, *Astyanacinus,* and *Ctenobrycon* from Colombia generated in this study for phylogenetic analyses.*Type locality.^a^[15].

| Species | Sample ID/Name | Locality /Basin | *rag2* | *COI* |
| --- | --- | --- | --- | --- |
| *Astyanax microlepis* | 7927 | Piedra Moler - Valle del Cauca*/upper Cauca | OR992015 | OR981620 |
| *Astyanax microlepis* | 7965A | Piedra Moler - Valle del Cauca*/upper Cauca | OR992016 | OR981621 |
| *Astyanax microlepis* | 7965B | Piedra Moler - Valle del Cauca*/upper Cauca | OR992017 | OR981622 |
| *Astyanax microlepis* | 7965C | Piedra Moler - Valle del Cauca*/ upper Cauca | OR992018 | OR981623 |
| *Astyanax* sp | UN636 | Santa Fé de Antioquia- Bolombolo/middle Cauca | OR992030 | OR981635 |
| *Astyanax* sp | A2907 | Ituango reservoir tail/middle Cauca | OR992019 | OR981624 |
| *Astyanax* sp | UN1556 | Puente Tarazá/lower Cauca | OR992020 | OR981625 |
| *Astyanax* sp | UN1557 | Puente Tarazá/lower Cauca | OR992022 | OR981627 |
| *Astyanax* sp | UN1565 | Puente Tarazá/lower Cauca | OR992031 | OR981636 |
| *Astyanax* sp | UN1560 | Puente Tarazá/lower Cauca | OR992023 | OR981628 |
| *Astyanax* sp | UN1564 | Puente Tarazá/lower Cauca | OR992029 | OR981634 |
| *Astyanax* sp | UN1561 | Puente Tarazá/lower Cauca | OR992021 | OR981626 |
| *Astyanax* sp | UN1566 | Puente Tarazá/lower Cauca | OR992024 | OR981629 |
| *Astyanax* sp | C37450 | Margento/lower Cauca | OR992025 | OR981630 |
| *Astyanax* sp | C11854 | Tres Cruces/lower Cauca | OR992026 | OR981631 |
| *Astyanax* sp | C11862 | Tres Cruces/lower Cauca | OR992027 | OR981632 |
| *Astyanax* sp | C11877 | Tres Cruces/lower Cauca | OR992028 | OR981633 |
| *Astyanax* sp | IAvH-CT-24548 | Barrancabermeja - Santander/middle Magdalena | - | OR064742 |
| *Astyanax* sp | IAvH-CT-24679 | Cimitarra - Santander/middle Magdalena | OR992033 | OR064743 |
| *Astyanax* sp | IAvH-CT-17785 | Colosó - Sucre/lower Magdalena | OR992032 | OR064741 |
| *Astyanacinus yarigues* | IAvH-23068 | Carmen de Chucurí -Santander/middle Magdalena | OR992034 | OR981637 |
| *Ctenobrycon caucanus* | UN1536 | Tarazá River*/middle Cauca | OR992035 | OR064739<sup>a</sup> |

**Table 2.**
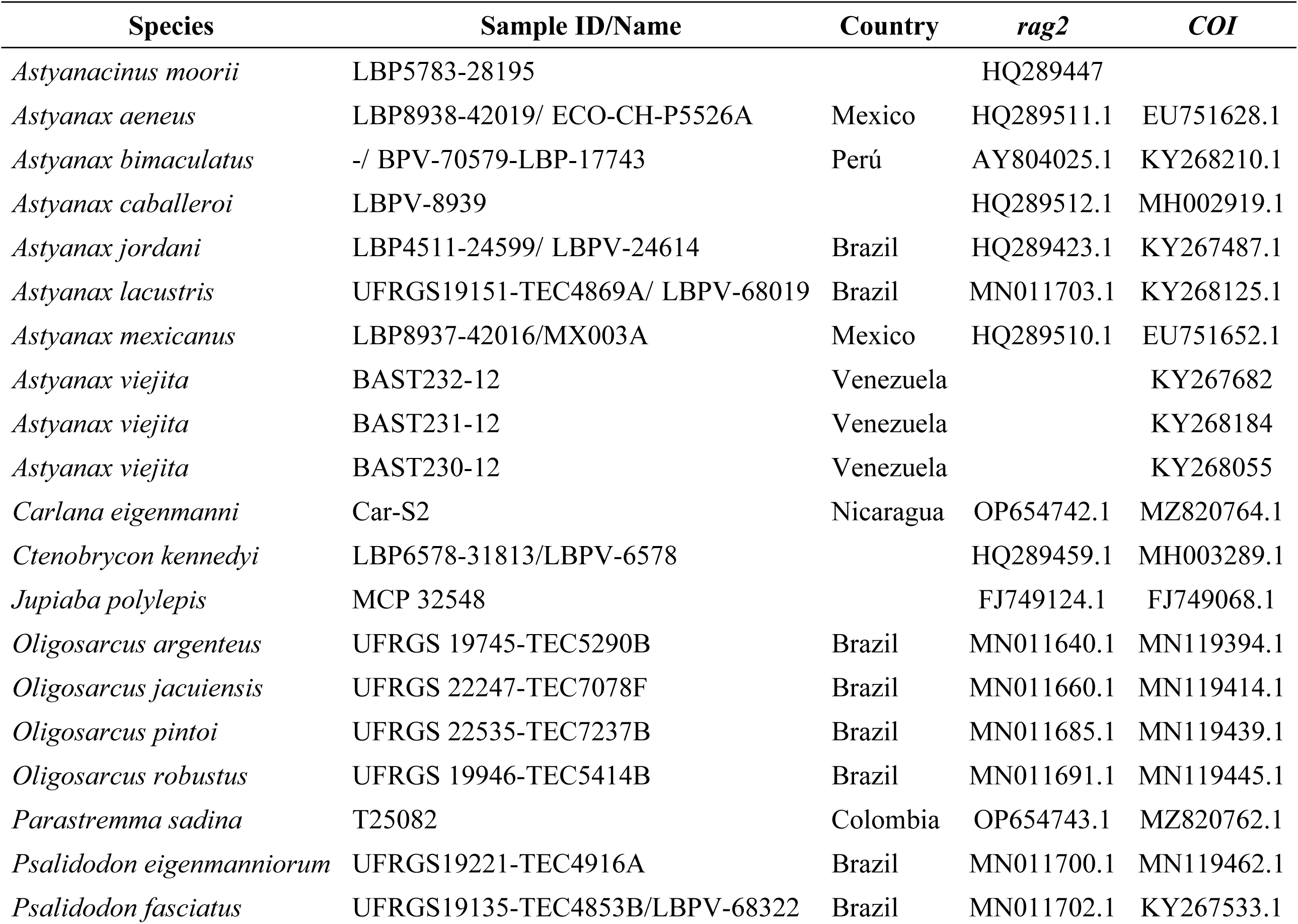
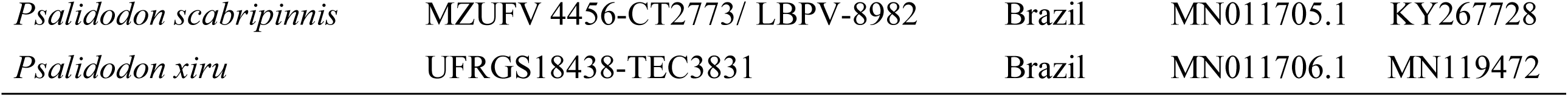
List of sequences retrieved from GeneBank for phylogenetic analysis.

Phylogenetic reconstruction was conducted using the ‘A la Carte’ workflow integrated in NGPhylogeny.fr [52]. Sequences were aligned with MAFFT [53] and the resulting alignment was optimized using TrimAl [54] to remove poorly aligned regions and gaps. The best-fit evolutionary model was determined using IQ-TREE [55], which identified TIM2e+I+R2 as the optimal model for the dataset based on the Bayesian Information Criterion (BIC). Bayesian Inference was performed in MrBayes v3.2 [56] implementing as selected evolutionary model, a General Time Reversible (GTR) model with a proportion of invariant sites and Gamma-distributed rate variation (+I+G; nst=6, rates=invgamma). The Metropolis-coupled Markov Chain Monte Carlo analysis consisted of two independent runs (nrun=2), each utilizing four chains (one cold and three heated), for 2,000,000 generations. To ensure scientific reproducibility, the analysis was executed with fixed starting seeds (Seed=5, Swapseed=5). Parameters and trees were sampled every 500 generations, and a 25% burn-in fraction was applied, discarding the initial 500,000 generations. Convergence and stationary distributions were assessed using Tracer v1.7.2 [57], ensuring that all parameters reached an Effective Sample Size (ESS) greater than 200. The consensus tree was rooted using *Jupiaba polylepis* as the outgroup and visualized using Newick Display [58] and iTOL [59].

### Geometric morphometrics

A total of 138 specimens were collected during the dry season. Each specimen was photographed in left lateral view (Fig 2), inside a diffuse light box, using a 12 MP Samsung digital camera, with focal length of 47 cm. Landmark digitization was conducted using the XYOM v. 3.0 application [60] (https://xyom.io/), marking on each image, eight type I landmarks (juxtaposition of tissues) and one type II landmark (tip of snout; [61]). The digitization process was repeated twice to ensure precision, and a Generalized Procrustes Analysis [62] was applied to align landmark coordinates (S2 Table), removing variation due to size, rotation, and position [63]. Body shape variables (partial warps and uniform components) were analyzed using linear discriminant analysis, implemented with the R packages MASS [64] for analysis, and ggplot2 [65] for visualization. Additionally, a classification approach based on cross-validation was applied using the R package caret [66] to evaluate model performance. The statistical significance of Euclidean distances to explore differences among genetic groups was assessed through 1,000 permutations using CLIC software [60]. Additionally, a multivariate regression was used to assess the effect of size on shape (allometry). When significant allometry was detected, size correction of shape was performed if the allometric slopes fitted a common model, which was evaluated using multivariate analysis of covariance (MANCOVA). A thin-plate spline graphical analysis was conducted using tpsRegr v.1.45 [67] and mean objects (residual coordinates obtained after Procrustes superimposition), generated with CLIC, to illustrate body shape deformations.

**Fig 2.**
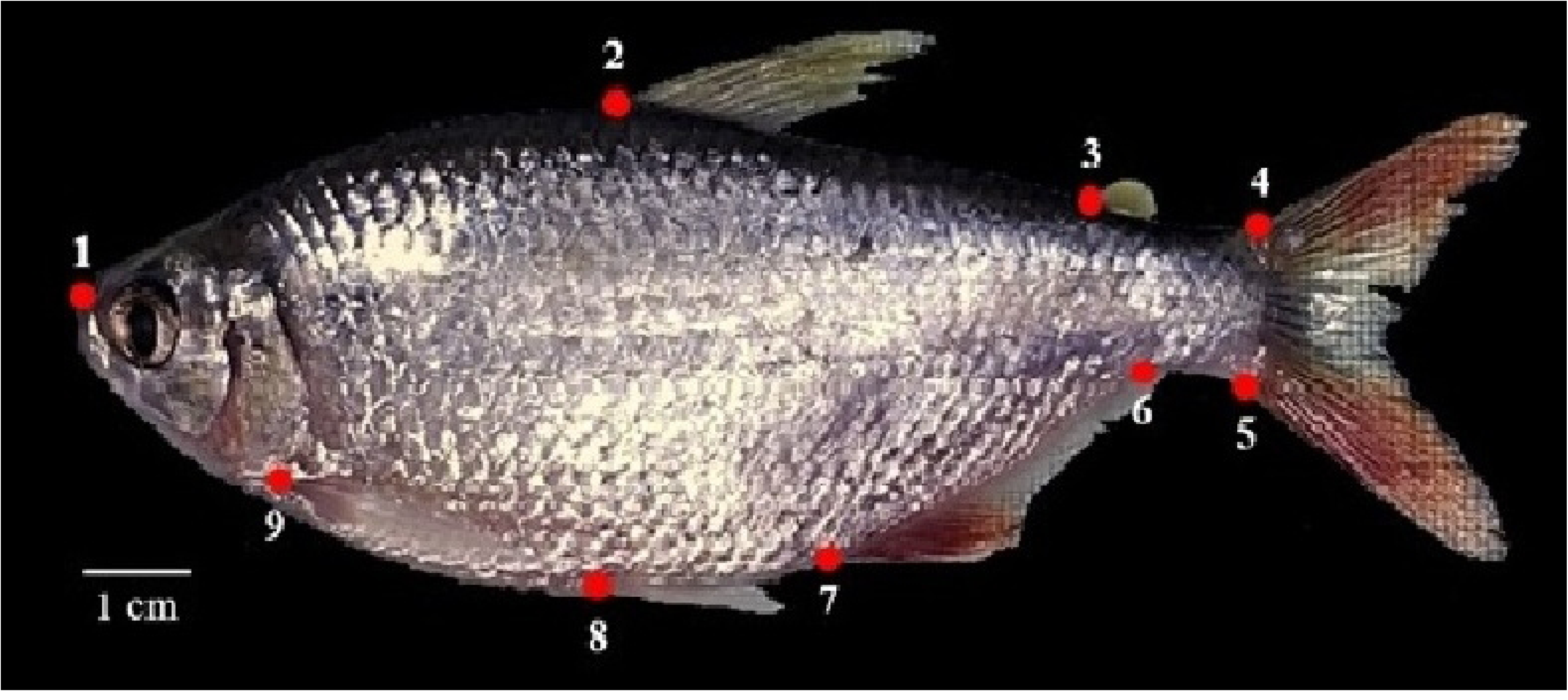
Landmarks selected as body shape coordinates for morphometric analysis of *Astyanax* species. 1, tip of snout; 2, dorsal-fin origin; 3, adipose-fin origin; 4-5, origin of uppermost and lowermost caudal-fin rays; 6-7, anal-fin insertion; 8, pelvic-fin origin; 9, pectoral-fin origin.

## Results

### STR development and polymorphism

Among the initially 42 evaluated loci, 17 were polymorphic, consistently amplified, and exhibited PIC values above 0.500 (Table 3). The number of alleles per locus ranged from 4 to 31, with an allelic range (Ra) of 105 to 381, and expected heterozygosity values between 0.656 and 0.965. Some of these loci exhibited a high fixation index, and 11 showed significant deviation from Hardy-Weinberg equilibrium. Due to limited sample quantity, loci Afa18, Afa36, and Afa38 were not amplified in the broader sample set. Consequently, a final suite of 14 loci was selected for subsequent population genetic analyses.

**Table 3.**
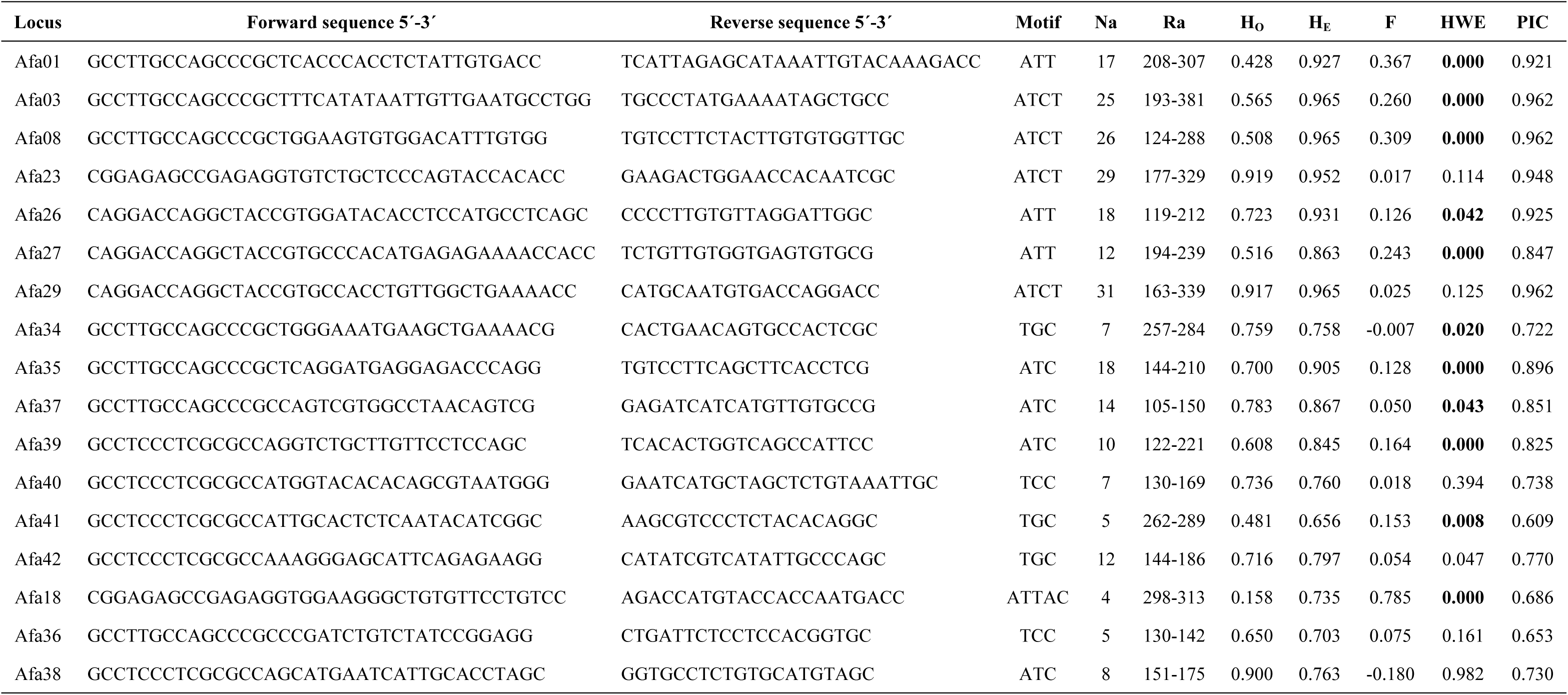
Characteristics of 17 loci isolated from *Astyanax* sp. Na: average number of alleles, Ra: allelic range, H_O_: observed heterozygosity, H_E_: expected heterozygosity, HWE: p-value of the Hardy-Weinberg equilibrium test, PIC: polymorphic information content. Bold values are significant in HWE. Universal Primers. A: GCCTCCCTCGCGCCA, B: GCCTTGCCAGCCCGC, D: CGGAGAGCCGAGAGGTG [38].

### Population genetics analysis

The Bayesian clustering analysis (K = 2) identified two distinct genetic stocks, hereafter referred to as Asp1 and Asp2 (Fig 3; S3 Table). Asp1 was the predominant stock in sectors S1–S4 and PHI, whereas Asp2 prevailed in the downstream sections S5, S6, and S7/S8.

**Fig 3.**
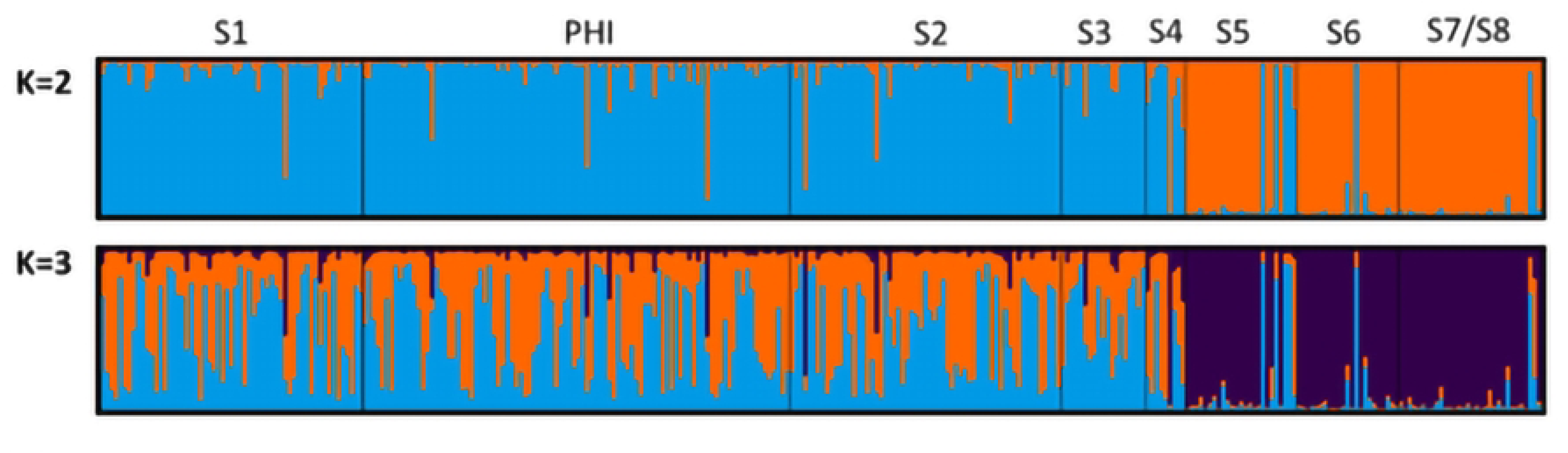
Histogram of genetic co-ancestry for K=2 (A) and K=3 (B) populations. Each vertical bar represents an individual, with color indicating the membership probability for each genetic stock (color). The K=2 model was supported by ΔK method, while K=3 was suggested by the MedMeaK, MaxMeaK, MedMedK and MaxMedK estimators. In both plots, blue and orange represent Stock 1 (Asp1) and Stock2 (Asp2), respectively.

Genetic differentiation was further supported by the AMOVA [F’_ST (7, 647)_ = 0.073, P= 0.001], pairwise structure indices (Table 4), and the DAPC (Fig 4), which showed a clear separation between downstream sectors (S5-S8) and both upstream (S1 and PHI) and dam-proximate sectors (S2 and S3). Furthermore, differentiation was higher and consistently significant across both estimators (F’_ST_ > 12 % and D_EST_ > 8%) when comparing the upstream and dam-proximate sectors (S1, PHI, S2, and S3) with the most downstream sites (S5, S6 and S7/S8).

**Fig 4.**
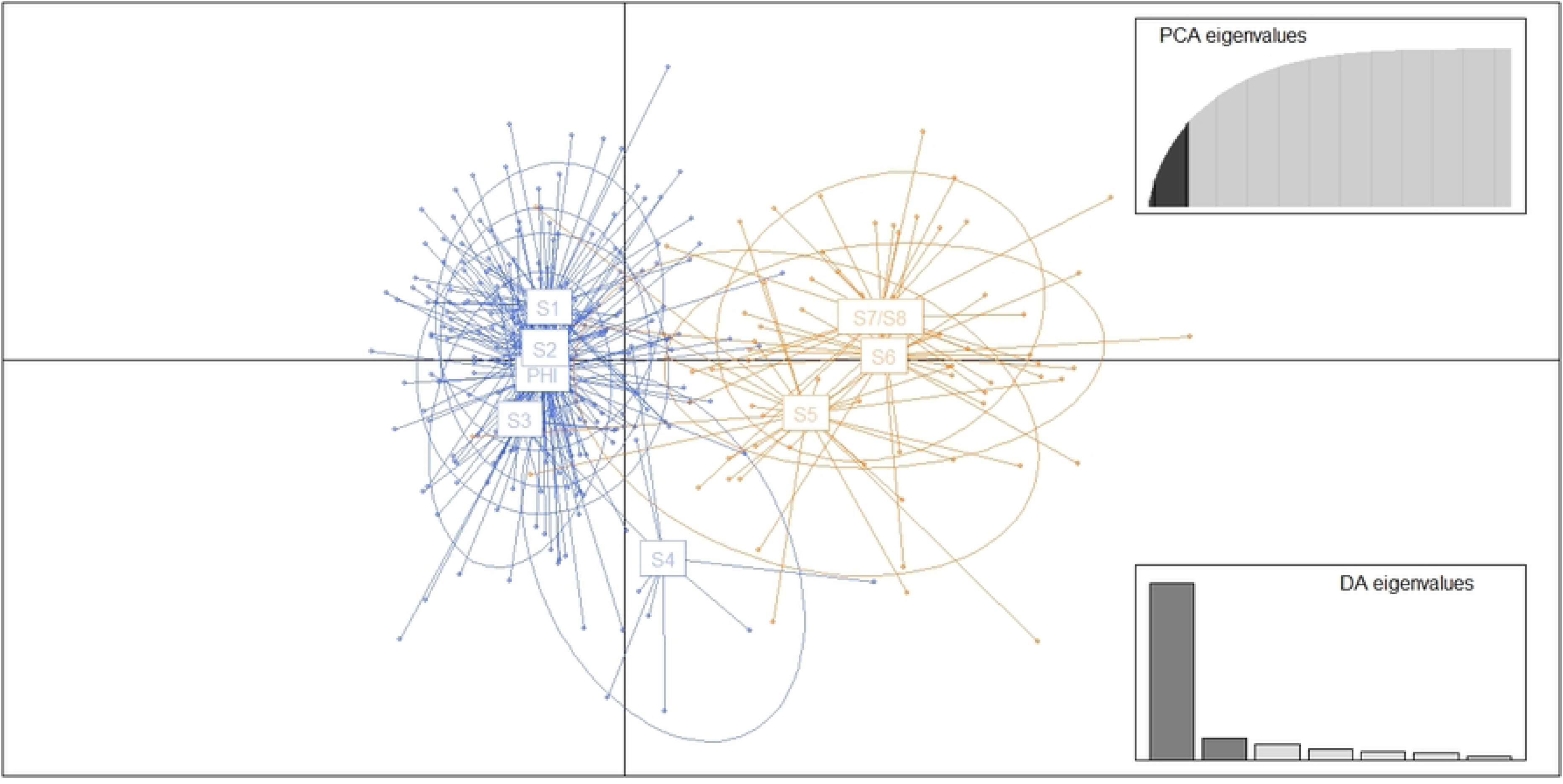
Discriminant analysis of principal components of samples of *Astyanax* sp.

**Table 4.** Pairwise indices of genetic structure of *Astyanax* sp. F’_ST_ below and D_EST_ above. Significant values in bold.

|  | S1 | PHI | S2 | S3 | S4 | S5 | S6 | S7/S8 |
| --- | --- | --- | --- | --- | --- | --- | --- | --- |
| S1 |  | -0.005 | 0.004 | 0.030 | 0.094 | **0.100** | **0.140** | **0.146** |
| PHI | -0.005 |  | -0.006 | 0.000 | 0.042 | **0.082** | **0.135** | **0.137** |
| S2 | -0.007 | 0.005 |  | 0.022 | 0.084 | **0.085** | **0.129** | **0.136** |
| S3 | 0.006 | 0.048 | 0.034 |  | 0.075 | 0.072 | **0.141** | **0.166** |
| S4 | 0.053 | 0.114 | 0.096 | 0.088 |  | 0.054 | 0.060 | 0.112 |
| S5 | **0.138** | **0.146** | **0.122** | 0.134 | 0.095 |  | 0.004 | 0.015 |
| S6 | **0.157** | **0.156** | **0.141** | **0.174** | 0.069 | 0.008 |  | -0.013 |
| S7/S8 | **0.165** | **0.165** | **0.150** | **0.195** | 0.123 | 0.011 | -0.014 |  |

Across the eight evaluated sectors (Table 5), *Astyanax* sp. exhibited comparable levels of genetic diversity (Ar: 7.7–8.7; H_O_: 0.549–0.694; H_E_: 0.827–0.855). However, a generalized heterozygote deficit was observed, with significant inbreeding coefficients (F_IS_ > 0.10) that were relatively higher in downstream areas. Consistent with these sector-wide patterns, the downstream stock (Asp2) displayed a higher inbreeding coefficient than its upstream counterpart, despite maintaining similar overall levels of genetic diversity.

**Table 5.**
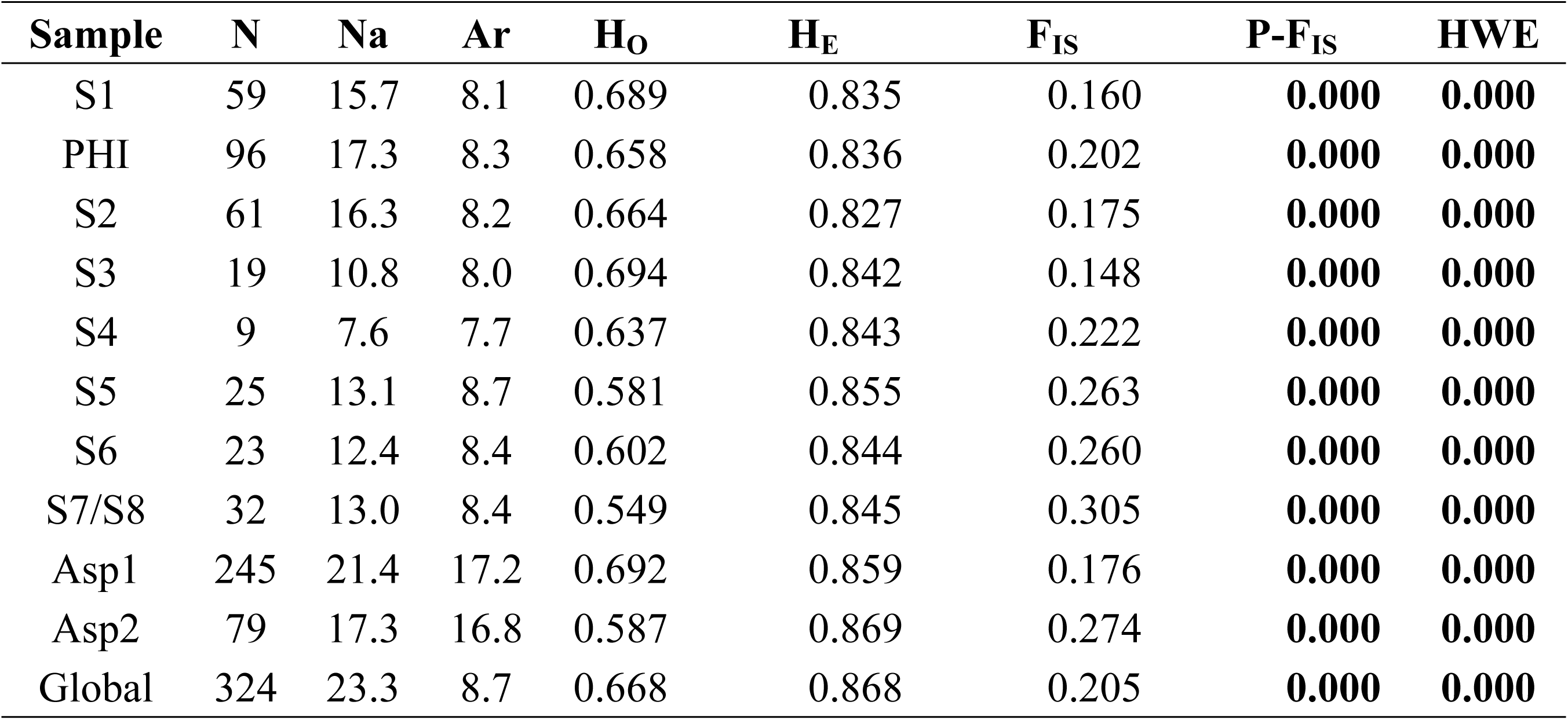
Genetic diversity and inbreeding coefficient in samples of *Astyanax* sp. N: sample size, Na: average number of alleles, Ar: allelic richness, H_O_: observed heterozygosity, H_E_: expected heterozygosity, F_IS_: inbreeding coefficient, P-F_IS_: p-value of the inbreeding coefficient, HWE: p-value of the Hardy-Weinberg equilibrium test. Bold values are significant. Asp1: Stock 1; Asp2: Stock2.

Evidence of recent bottleneck events was observed across all sectors and stocks, with significant results obtained from both estimation methods in at least one of the three mutational models (Table 6). Successful estimations of effective population size (Ne) were limited; however, values exceeding 600 were recorded for sectors in the vicinity of the dam.

**Table 6.** Bottleneck and effective population size estimations of *Astyanax* sp. N: sample size, IAM, TPM, and SMM: p-values from the method used by BOTTLENECK v. 1.2.02 [45] for the three mutational models, M: index proportion between number of alleles and allelic range, indicating recent bottleneck events when less than 0.68 [46], N_e_: estimated effective population size, using the linkage disequilibrium method, IC: Jackknife confidence intervals. Inf: infinite estimation. Asp1: Stock 1; Asp2: Stock2.

| Sample | N | M ratio | IAM | TPM | SMM | $N_e$ | IC |
| --- | --- | --- | --- | --- | --- | --- | --- |
| S1 | 59 | 0.254 | **0.000** | **0.003** | 0.852 | 668.5 | 248.7-inf |
| PHI | 96 | 0.270 | **0.000** | **0.015** | 0.966 | 2240 | 536.3-inf |
| S2 | 61 | 0.277 | **0.000** | **0.039** | 0.988 | 1138.9 | 405.2-inf |
| S3 | 19 | 0.221 | **0.000** | **0.000** | 0.729 | inf | 100.7-inf |
| S4 | 9 | 0.213 | **0.000** | **0.001** | 0.077 | inf | 32.5-inf |
| S5 | 25 | 0.236 | **0.000** | **0.002** | 0.108 | inf | 120-inf |
| S6 | 23 | 0.249 | **0.000** | **0.010** | 0.687 | inf | 144.6-inf |
| S7/S8 | 32 | 0.267 | **0.000** | **0.000** | 0.195 | inf | 166.2-inf |
| Asp1 | 245 | 0.277 | **0.000** | **0.024** | 1.000 | inf | 2093.4-inf |
| Asp2 | 79 | 0.257 | **0.000** | **0.000** | 0.380 | inf | 2050.6-inf |
| Global | 324 | 0.280 | 0.000 | **0.012** | 0.997 | 1364.0 | 850.8-3150.1 |

The overall population displayed an Ne estimate surpassing 1000, with a 95% confidence interval ranging from 850 to 3150.

### Phylogenetics analysis

The phylogenetic tree reveals two well-supported divergent clades encompassing the samples from the middle and lower Cauca River (Fig 5). Lineage 1 (blue clade) clustered samples corresponding to Asp1 (Fig 3), alongside sequences of *Astyanax microlepis* and *A. viejita*. Conversely, Lineage 2 (orange clade) grouped samples corresponding to Asp2 (Fig 3) with samples of *Astyanax* sp. from the Magdalena River. Both clades were distinct from all other reference sequences. Furthermore, previous taxonomic assignment to *Psalidodon fasciatus* is unequivocally dismissed, as *Psalidodon* samples occupy a distinct phylogenetic position.

**Fig 5.**
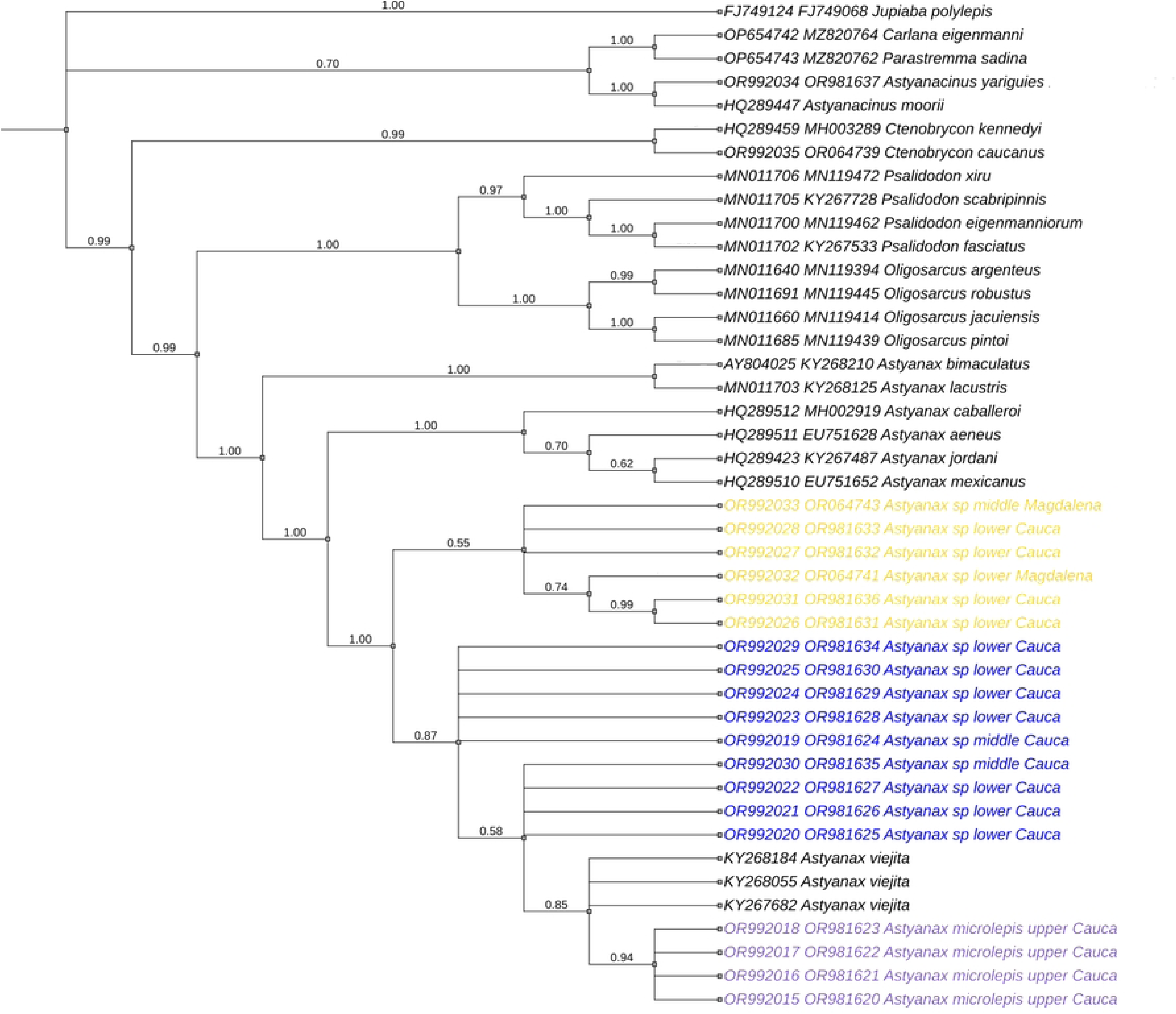
Bayesian phylogenetic tree based on concatenated mitochondrial (*COI*) and nuclear (*rag2*) gene sequences, illustrating the relationships of *Astyanax* sp. from the Cauca River. Blue and orange clades correspond to stocks Asp1 and Asp2, respectively, as identified in the genetic structure analyses (Fig. 3). The purple clade represents *A. microlepis* collected from its type locality in the upper Cauca River. The analysis incorporates GenBank sequences of *A. viejita* [4] from the Lake Maracaibo basin and other congeners (Table 2). Members of the genera *Astyanacinus*, *Carlana*, *Ctenobrycon*, *Jupiaba*, *Oligosarcus*, *Parastremma*, and *Psalidodon* were utilized as outgroups.

### Geometric morphometric analyses

Linear discriminant analysis revealed clear morphological differentiation among three analyzed groups of *Astyanax*, based on body shape variables (Fig 6). The first discriminant function accounted for 58.12% of the total variance, separating *A. microlepis* (red triangles) from the other groups. The second discriminant function accounted for 41.88% of the variance, separating *Astyanax* sp. 1 (Asp1; blue squares) from *Astyanax* sp. 2 (Asp2; orange circles).

**Fig 6.**
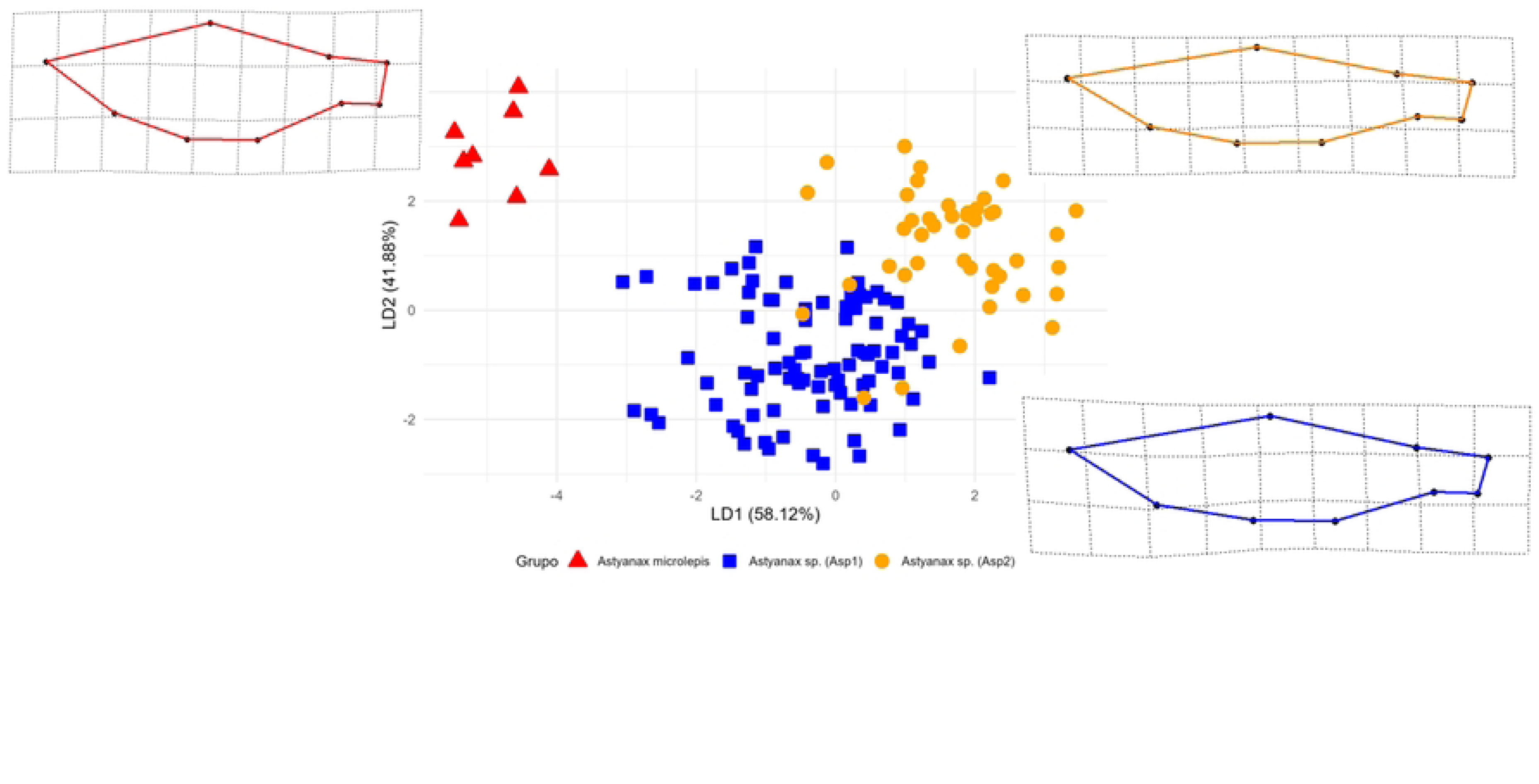
Linear discriminant analysis plot showing morphological differentiation among three *Astyanax* groups, based on body shape variables (partial warps and uniform components). The first two linear discriminants (LD1 and LD2) explain 58.12% and 41.88% of the total variation, respectively. Red triangles: *Astyanax microlepis*; blue squares: *Astyanax* sp. 1 (Asp1); orange circles: *Astyanax* sp. 2 (Asp2). Thin-plate spline deformation grids reflect the relative deformation from the consensus shape, highlighting intergroup variation.

Thin-plate spline deformation grids based on mean shapes revealed distinct morphological patterns among groups (Fig 7). *Astyanax microlepis* exhibited a more robust body profile, Asp1 displayed an intermediate and slightly deeper body shape, while Asp2 was characterized by a more streamlined morphology. Specifically, Asp1 exhibited a shorter head than Asp2, a more posterior position of the adipose fin, and a more ventral position of both anal-fin insertion and pelvic-fin origin.

**Fig 7.**
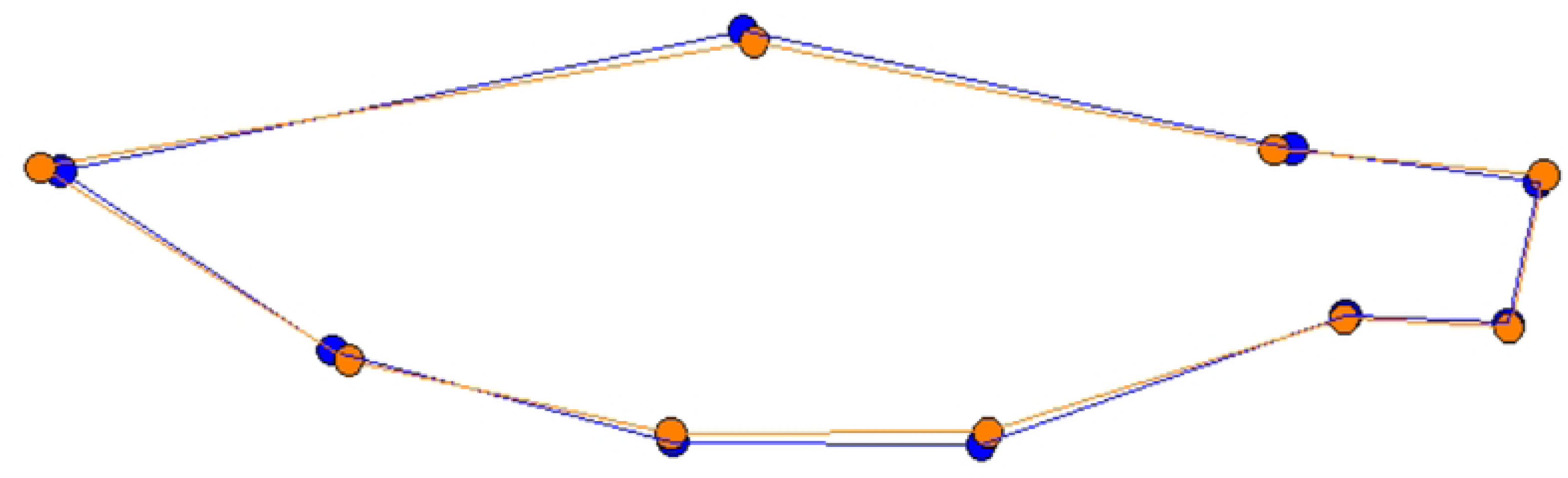
Body shape average of genetic groups of *Astyanax* sp. after generalized Procrustes analysis

Morphological differences among groups were supported by both allometric and non-allometric components of body shape variation, quantified using Euclidean distances and tested for significance through 1,000 permutations (Table 7).

**Table 7.** Euclidean distances uncorrected (ED) and size-corrected (cED), along with their statistical significance (P), for body shape differences among *Astyanax* groups.

| Comparison | ED | P(ED) | cED | P(cED) |
| --- | --- | --- | --- | --- |
| *Astyanax microlepis* vs. *Astyanax* sp. 1 (Asp1) | 0.133 | 0.000 | 0.133 | 0.000 |
| *Astyanax microlepis* vs. *Astyanax* sp. 2 (Asp2) | 0.144 | 0.000 | 0.143 | 0.000 |
| Asp1 vs. Asp2 | 0.027 | 0.002 | 0.025 | 0.000 |

The classification approach, based on cross-validation, showed high classification accuracy among the three groups. *Astyanax microlepis* specimens were correctly classified in 100% of cases (8/8), Asp1 in 98.85% (86/87), and Asp2 in 88.37% (38/43). The overall classification success rate was 95.65%, indicating high morphological differentiation among groups (Table 8).

**Table 8.** Cross-validation classification results, based on body shape variables among *Astyanax* groups.

|  | *A. microlepis* | Asp1 | Asp2 | % |
| --- | --- | --- | --- | --- |
| *Astyanax microlepis* | 8 | 0 | 0 | 100 |
| *Astyanax* sp. 1 (Asp1) | 0 | 86 | 1 | 98.85 |
| *Astyanax* sp. 2 (Asp2) | 0 | 5 | 38 | 88.37 |
| **Total** |  |  |  | 95.65 |

## Discussion

Neotropical freshwater fish evolution reflects a complex interplay between contemporary demographic processes and deep-time cladogenesis. In this study, we integrated newly developed microsatellite loci to disentangle these scales in *Astyanax* sp. from the Cauca River. Although microsatellites are established tools for testing microevolutionary hypotheses [37; 68, 69], this work represents only the second instance in Colombia where species-specific markers have been successfully developed for this group, following the previous effort on *Ctenobrycon caucanus* (formerly *Astyanax caucanus*) [15]. The high-resolution of these new loci enabled the detection of fine-scale genetic discontinuities in the middle and lower sections of the basin, and provided highly informative baselines for monitoring the genetic health of *Astyanax* populations. This is particularly relevant in sectors impacted by the Ituango dam, where microevolutionary forces are actively shaping the genomic landscape.

Genetic diversity in terms of alleles and expected heterozygosity was similar and high between both genetic stocks, compared to other Neotropical characiforms (Na: 10.1, H_E_: 0.675;[70]) and cis-Andean congeners (e.g. *Astyanax scabripinnis*) [12]. Nonetheless, both genetic stocks showed high inbreeding levels, particularly higher in Asp2 from the lower section of the river. These levels of genetic diversity and inbreeding were comparable to those reported for *Ctenobrycon caucanus* (H_E_: 0.832-0.840; F_IS_: 0.307-0.351; [15]), another sympatric species in the Cauca River, and to Central American congeners such as *A. aeneus* and *A. caballeroi* [14].

Although much is still unknown about the life cycle of Acestrorhamphidae species from the Magdalena basin, similar genetic features among coinhabiting species, like *Astyanax* sp. And *C. caucanus*, such as inbreeding, genetic diversity, and bottleneck events (see [71]), may reflect a shared biogeographic history, coupled with as-yet-unknown intrinsic biological factors of their life cycle.

Despite the observed genetic evidence indicating bottleneck events in the samples analyzed, the estimated effective population size for Asp1 was high, consistent with reports for other species of the Magdalena basin [71]. This is also congruent with abundant populations [72–74], as observed in *Astyanax* sp. and *A. microlepis*, reported to be the most common species, in terms of abundance and biomass in the basin, even in dammed environments [9]. Such demographic robustness suggests that while these populations experience historical or contemporary fluctuations, they maintain large reproductive pools capable of preserving high levels of genetic variation.

The implications of these population-level findings extend beyond contemporary demographics. To determine whether the Asp1 and Asp2 genetic stocks represent incipient speciation or established evolutionary lineages, we bridged the gap between micro- and macroevolutionary scales by integrating a dual-gene phylogenetic approach (*COI* and *rag2*) with geometric morphometrics. This framework allowed us to assess the taxonomic congruence of the clusters detected by microsatellites in a broader evolutionary context, providing a robust characterization of the diversity within trans-Andean *Astyanax*. Collectively, these independent lines of evidence are essential for a more comprehensive understanding of the complex biodiversity of trans-Andean ichthyofauna.

Such integration is critical as taxonomic uncertainties at the species level, together with the non-monophyletic condition defining the *sensu lato* concept of *Astyanax* (still in use), have led to the conspicuous proliferation of samples identified only to the genus level in ichthyological collections [4,5]. This has also been the case for *Astyanax* sp. from the Magdalena basin in Colombia. This species had been traditionally identified as *Astyanax fasciatus* [7], until this nominal species was reallocated in the recently resurrected *Psalidodon*, along with evidence indicating that known species of this genus are geographically restricted to Atlantic basins of Brazil, as the Rio São Francisco (type locality of *P. fasciatus*) and to the Paraná-Paraguay basin [2,5]. This taxonomic uncertainty of the Magdalena species raises concerns for its populations from the Cauca River, given their trophic importance into the local fish assemblages, reports of increasing human consumption, and environmental threats in the basin, by pollution of aquatic habitats and by habitat fragmentation due to the Ituango dam [9,35,75].

Phylogenetic analyses showed that the two genetic stocks identified by microsatellites also fall into two distinct lineages strongly supported using both mitochondrial and nuclear genes. Lineage 1 consisted of samples from Asp1, but surprisingly, this lineage also encompasses samples of *A. microlepis* (from its type locality, in the upper basin of the Cauca River), and *A. viejita* (from the Maracaibo Lake basin).

*Astyanax microlepis* has been reported throughout the Cauca River [10,76] and was recovered as sister to the clades *A. bimaculatus* and *A. argentatus* [5]. Conversely, sequences of *Astyanax viejita* are exclusive from the Maracaibo Lake basin in Venezuela [3], with no previous reports of its presence in Colombia [6]. Lineage 2 comprised samples from Asp2, including sequences of *Astyanax* sp. collected in the main channel of the middle section of the Magdalena River.

Despite genetic similarity between *Astyanax microlepis* and Asp1 (based at least on partial *cox1* sequences), results of the linear discriminant analysis revealed clear morphological differentiation between both species. Sympatric divergence in *Astyanax* has been documented in the Nicaraguan lakes Managua [comprising the deep-bodied *A. nasutus* Meek, 1907 and *A. nicaraguensis* Eigenmann & Ogle, 1907, and the elongate *A. bransfordii* (Gill, 1877)] and Nicaragua (where *A. nicaraguensis* and *A. bransfordii* also co-occur). In both lakes, these species exhibit differences in premaxilla shape, in addition to differences in body configuration. Such variations correlate strongly with trophic ecology, suggesting adaptive divergence [22,73]. Similar findings have been reported for *A. aeneus* and *A. caballeroi* in Lake Catemaco, Mexico [14,19].

This discordance between phenotypic and genotypic variation can be explained by some non-mutually exclusive hypotheses. One possible explanation is that morphotypes are undergoing incipient speciation. In this scenario, the lack of widespread genetic differentiation is an expected consequence of recent divergence, which leads to incomplete sorting of ancestral lineages [77–79]. Furthermore, a divergence-with-gene-flow model may explain how strong selection on a few “genomic islands” can still create distinct phenotypes under these conditions [79,80]. Our results are concordant with this model, which is supported by examples in related species like *A. aeneus* and *A. caballeroi* that maintain distinctiveness amid significant admixture [14].

The discordance observed between phenotypic and mitochondrial data for *A. microlepis* and Asp1 can be explained by a scenario of allopatric divergence followed by secondary contact. This evolutionary model has been previously invoked to explain the lack of correspondence between pronounced morphological divergence and genetic structure in sympatric *Astyanax* morphotypes [19]. In such cases, the maintenance of distinct phenotypes despite gene flow depends on the strength of natural and sexual selection, as well as the robustness of reproductive isolating barriers [19, 81]. Under this scenario, our findings suggest a process of unidirectional introgression, where recent hybridization has eroded the molecular signals of previous isolation at the organellar level. Similar patterns of secondary contact and subsequent genetic exchange have been documented in other Neotropical contexts; for instance, the complex hydrological history of the San Juan River basin seems to reflect secondary contact between deep- and elongated-bodied *Astyanax* morphs [22], while significant introgression has been reported in *Herichthys*, following recent contact between divergent lineages [82]. Taken together, these precedents suggest that the phenotypic integrity of *A. microlepis* and Asp1 is likely maintained by strong divergent selection, even if their mitochondrial lineages have been homogenized. However, as mitochondrial DNA represents only a single maternal locus, further research using high-resolution genomic tools is required to confirm the extent, directionality, and nuclear impact of this gene flow.

A third possibility is that these morphotypes represent a case of stable polymorphism within a single species. Under this model, divergent morphs develop from a single gene pool, where the developmental trajectory is maintained by balancing selection or frequency-dependent mechanisms rather than a transitional phase towards speciation [82–85]. This scenario has been strongly suggested for other *Astyanax* populations, where marked morphological divergence occurs in the absence of genetic structure, such as in Lake Catemaco [14] and the San Juan River basin [22]. Similar examples of stable trophic polymorphism include the cichlid *Herichthys minckleyi* [82] and the mouth asymmetry in *Perissodus microlepis* [86].

Closely related to this is the hypothesis of phenotypic plasticity, where a single genotype produces alternative phenotypes in response to environmental cues. In Neotropical fishes, plasticity is frequently invoked to explain ecomorphological divergence in the absence of genetic differentiation. For instance, some authors suggested that the extreme trophic morphotypes in Mesoamerican *Astyanax* (formerly *Bramocharax*) are plastic expressions within a single lineage [17]. Similarly, the body shape in *Astyanax* and *Psalidodon* has been shown to be directly influenced by lentic-lotic gradients of their habitats [23].

However, both hypotheses (stable polymorphism and phenotypic plasticity) are difficult to reconcile with the geographical and ecological patterns observed in the species of our study. First, geographic distributions are discordant: while Asp1 is widespread throughout the Magdalena and Cauca river drainages, *A. microlepis* is notably absent in the Magdalena River [6,76]. If these forms were merely plastic responses or a localized polymorphism, similar phenotypic ranges would be expected in the whole Magdalena basin. Second, and most importantly, both phenotypes occur syntopically across all sampled habitats in the Cauca River (including lotic, lentic, and artificial reservoir environments; [76]; this study). The fact that *A. microlepis* and Asp1 maintain their distinct morphologies, despite sharing the same range of habitats suggests a significant heritable component, rather than purely environmentally induced morphs.

In summary, while the mitochondrial genome shows lack of differentiation (potentially due to recent introgression or a selective sweep), the persistence of these phenotypes in syntopy points toward a stable evolutionary divergence. Therefore, *A. microlepis* and Asp1 likely represent separate evolutionary lineages with divergence occurring at the nuclear level, a pattern that requires confirmation through high-resolution genomic tools to fully unravel the history of this complex group.

Conversely, the more genetically divergent pair of Asp1 and Asp2 displayed greater overall morphological similarity. Despite their general resemblance, our analysis detected fine-scale yet significant differences, particularly in head length, position of pectoral and adipose fins, and displacement of the origin of the dorsal and anal-fins. This phenomenon, where lineages diverge genetically while their morphology remains constrained, points toward non-mutually exclusive evolutionary mechanisms.

An explanation for the conserved morphology in the face of genetic drift is the action of strong stabilizing selection, coupled with niche conservatism [87]. If both lineages inhabit similar ecological contexts, as their sympatric distribution in the Cauca River basin suggests, any significant deviation from an optimal, time-tested body plan would likely be disadvantageous and selected against. Consequently, while neutral genetic markers accumulate differences over time, functional morphology of organisms remains evolutionarily conserved by selection. This process results in the formation of cryptic species, which are genetically distinct but physically almost identical, a pattern increasingly uncovered in diverse taxonomic groups, including Neotropical freshwater fishes (e.g. [88–90]).

Alternatively, the subtle yet significant differences observed in head profile and relative position of fin insertions between Asp1 and Asp2, may reflect divergent evolutionary trajectories resulting from their independent histories, rather than specific adaptations to micro-niches within their shared habitat. Common inherited genetic architecture may impose developmental limitations to respond to selective pressures similarly [91]. Therefore, the morphological similarity between these stocks suggests a conserved body plan, potentially maintained by stabilizing selection or representing a symplesiomorphic condition within the clade. Distinguishing between these scenarios (the retention of ancestral traits [92] versus parallel evolution toward a common optimum [93]) will require broader morphometric analyses. Integrating additional species from the same clade into a phylomorphospace framework would clarify whether this shared morphology is a derived convergence or reflects the ancestral pattern of the group.

In conclusion, integrated molecular and morphometric approaches provide compelling evidence for the existence of two distinct evolutionary lineages within *Astyanax* sp. in the Cauca River. Additionally, the grouping of *A. microlepis*, *A. viejita*, and Asp1 into a single mitochondrial lineage, despite their differing geographic distribution and morphological distinctness, introduces a novel hypothesis regarding the taxonomic status and evolutionary history of these species. This study not only challenges traditional taxonomic boundaries but also provides a crucial molecular foundation for future systematic revisions. Ultimately, recognizing these lineages is essential for the design of effective management and conservation strategies, aimed at preserving the ichthyofaunal genetic diversity in the Magdalena and Maracaibo basins.

## Supporting information

S1 Table. Microsatellite loci and primer pairs developed and tested for *Astyanax* sp. in this study. Universal Primers. A: GCCTCCCTCGCGCCA, B: GCCTTGCCAGCCCGC, D: CGGAGAGCCGAGAGGTG [38].

S2 Table. Data matrix for the geometric morphometric analysis of *Astyanax* species. The table provides the unique specimen identifier (Sample ID), the assigned genetic lineage or group (Genetic stock), and the two-dimensional raw coordinates (x, y) derived from the anatomical landmarks digitized for body shape evaluation.

S3 Table. Multilocus genotypes of 14 microsatellite loci for *Astyanax* sp. The first row contains summary metadata: number of loci, individuals, populations (genetic stocks), and sample sizes. The third row provides individual identifiers (Sample ID), sampling sections (genetic stocks), and locus-specific information (name and repeat motif).

